# Recurrent exon-deleting activating mutations in *AHR* act as drivers of urinary tract cancer

**DOI:** 10.1101/2021.11.09.468005

**Authors:** Judith M. Vlaar, Anouska Borgman, Eric Kalkhoven, Denise Westland, Nicolle Besselink, Charles Shale, Bishoy M. Faltas, Peter Priestley, Ewart Kuijk, Edwin Cuppen

**Author notes:** These authors contributed equally to this work: Ewart Kuijk, Edwin Cuppen. **Competing interests** The authors declare no competing interests.

## Abstract

Bladder cancer has a high recurrence rate and low survival of advanced stage patients. Few genetic drivers of bladder cancer have thus far been identified. We performed in-depth structural variant analysis on whole-genome sequencing data of 206 metastasized urinary tract cancers. In ~10% of the patients, we identified recurrent in-frame deletions of exons 8 and 9 in the aryl hydrocarbon receptor gene (*AHR*^*Δe8-9*^), which codes for a ligand-activated transcription factor. Pan-cancer analyses show that *AHR*^*Δe8-9*^ is highly specific to urinary tract cancer and mutually exclusive with other bladder cancer drivers. In *AHR*^*Δe8-9*^ the ligand-binding domain is disrupted and we show that this results in ligand-independent AHR-pathway activation. In bladder organoids, *AHR*^*Δe8-9*^ induces a transformed phenotype that is characterized by upregulation of AHR target genes, downregulation of differentiation markers and upregulation of genes associated with stemness and urothelial cancer. Furthermore, *AHRΔ*^*e8-9*^ expression results in anchorage independent growth of bladder organoids, indicating tumorigenic potential. DNA-binding deficient *AHR*^*Δe8-9*^ fails to induce transformation, suggesting a role for AHR target genes in the acquisition of the oncogenic phenotype. In conclusion, we show that *AHR*^*Δe8-9*^ is a novel driver of urinary tract cancer and that the AHR pathway could be an interesting therapeutic target.

## Introduction

Bladder cancer is the fourth most common cancer in men with a 5-year survival rate of about 77% ^1^. Despite risk stratification, the recurrence rate of bladder cancer is high and requires ongoing monitoring and treatment, which makes it also the costliest malignancy out of all cancers ^2^. With recent advances, the therapeutic options have been expanded to include immune checkpoint inhibitors and antibody-drug conjugates, while clinical studies for targeted drug approaches are ongoing ^3,4^. However, treatment and overall survival of cases with advanced stages of bladder cancer have not really improved in the last decades as compared to other cancers and there remains a need for the identification of drivers of bladder cancer tumorigenesis to effectively target early-stage disease and reduce tumor progression ^5,6^.

Genomic analyses of bladder cancer of various stages and grades have shown that bladder cancer is a heterogeneous disease of which the high mutation burden is largely driven by activation of APOBEC mutagenesis ^7–9^. Molecular characterization revealed activating *FGFR3* alterations in ~16% of patients, with a higher prevalence in non-muscle invasive bladder cancer ^9,10^. Furthermore, positive selection of mutated genes in the RTK-Ras-PI3K pathway (such as *FGFR3*, *PIK3CA*, and *ERBB2*), the p53-Rb pathway (such as *TP53*, *RB1*, and *ATM*), and genes involved in chromatin remodeling (*KDM6A* and *KMT2D*) has been identified in biopsies of patients with bladder cancer ^7,9^.

A recent study identified a rare non-synonymous mutation (Q383H) in *AHR* as an APOBEC-associated hotspot mutation in bladder cancer, suggesting *AHR* as a potential driver of bladder cancer ^11^. *AHR* is a ligand activated transcription factor involved in the cellular response to toxic aromatic hydrocarbons as well as cell type and context-specific physiologic functions ^12^. Increased *AHR* expression and activity have been associated with poor prognosis and survival for multiple cancers such as breast, lung, and upper urinary tract cancers ^13^. However, the relevance and functional consequences of cancer-associated *AHR* mutations in bladder cancer are still poorly understood.

Here, we describe the identification of novel recurrent exon-deleting *AHR* alterations in the pan-cancer whole-genome sequencing database of metastatic cancers from Hartwig Medical Foundation. Using state-of-the-art data analysis tools with improved functionality for detecting structural variants we identified a previously unnoticed recurrent in-frame deletion of exons 8 and 9 in *AHR*. Together with the Q383H point mutation and *AHR* gene amplifications, *AHR* variants comprise ~19% of patients with urinary tract cancer. We demonstrate that the recurrent mutations lead to constitutively activated AHR signaling and induces an oncogenic phenotype in bladder cells. Our results suggest that aberrant AHR signaling is an important driver of urothelial tumorigenesis.

## Results

### *AHR* is frequently mutated in urinary tract cancers

To identify genetic factors involved in urinary tract cancers, we analyzed the Hartwig database, which represents whole-genome sequencing data of solid metastatic tumors and normal tissue of 4500 patients ^14^. Utilizing improved structural variant detection algorithms ^15,16^, we detected a novel deletion spanning exons 8 and 9 of the *AHR* gene (*AHR*^*Δe8-9*^) in 22 (10.7%) out of the 206 urinary tract samples (Fig. 1a, 1b). The *AHR*^*Δe8-9*^ is identified in several urinary tract cancer subtypes, most of which are in bladder cancer (12/22) and pyelum (7/22) (Supplementary table 1). In the complete pan-cancer data set, the *AHR*^*Δe8-9*^ mutant is identified in only three non-urinary tract samples. The identified exonic deletion was validated in RNA-sequencing, which was available for 129 out of 206 urinary tract samples. Read support for the AHR exon 7-10 splice junction was identified in all urinary tract samples harboring *AHR*^*Δe8-9*^ (n = 16; Supplementary table 1) and confirmed the in-frame loss of exons 8 and 9 for six samples in which the genomic start position of the deletion is located within the coding sequence of exon 8.

**Figure 1.**
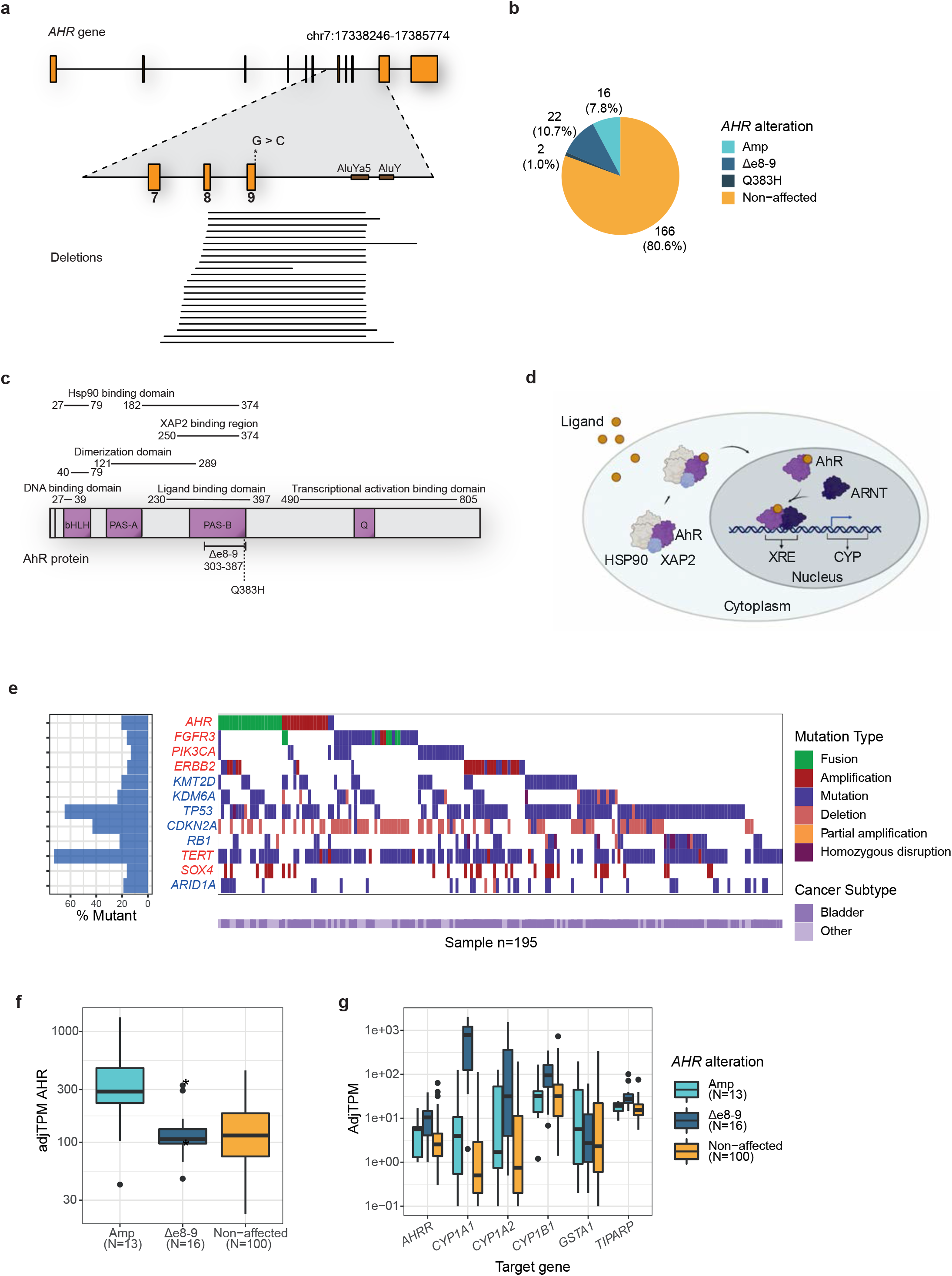
*AHR* alterations in somatic metastatic urinary tract patients. a. Schematic representation of *AHR* gene (ENST00000242057.4 - GRCh37) with a magnification of the genomic region around exon 7 - 9 (Chr7:17372389-17378532). The point mutation *Q383H* (c.1149G>C) is marked with an asterisk. The black bars in the lower part of the figure represent the extent of the genomic deletions identified in the urinary tract cancer patients. See supplementary table 2 for genomic positions of the deletions. b. Pie chart of the number of detected *AHR* alterations: *AHR* amplification (Amp), *AHR*^*Δe8-9*^, *AHR*^*Q383H*^ or non-affected samples in the urinary tract patient cohort of the Hartwig database. c. Schematic illustration of the AHhR functional domains and the positions of *AHR*^*Q383H*^ and *AHR*^*Δe8-9*^. The deletion (*AHR*^*Δe8-9*^) disrupts the PAS-B domain, ligand-binding domain (LBD), and XAP2 and HSP90 protein interaction domains. d. Canonical AHhR signaling pathway. (Figure created with BioRender.com). e. Visualization of top 12 affected genes per mutation type in urinary tract cancer samples of the Hartwig database ^14^. Gene names are colored for oncogenes (red) and tumor suppressor genes (blue). f. Boxplot of the adjusted TPM values for *AHR* transcript. Asterisks indicate two samples that contain the deletion (*AHR*^*Δe8-9*^) but also have an amplification of the *AHR* gene. g. Boxplot of the adjusted TPM values of AHR target genes.

The relatively small size of the *AHR* deletion (~ 3kb) may explain why it has not been identified in previous studies ^9,17^. Most of the 3’ breakpoint junctions are positioned in a narrow window of 30 bp between 2 *Alu* elements (*AluYa5* and *AluY*) and overlap with the 3’ site of the *AluYa5* element (Fig. 1a). Such inverted *Alu* repeats have been identified as strong initiators of genetic instability ^18^. The high prevalence in urinary tract cancer of such a rare event strongly suggests positive selection. Analysis of an independent RNA-sequencing dataset of urinary tract cancers from Weill Cornell Medicine confirmed the presence of the 7-10 splice junction in 8% (2/24) of samples. One of the bladder cancer samples with the detected splice junction was of primary origin (out of 8 primary tumors in this dataset). Together, these results demonstrate that the *AHR*^*Δe8-9*^ is present in primary and metastatic urinary tract cancer (Supplementary table 3).

To identify additional events in the *AHR* gene we performed a targeted analysis of the Hartwig database. We identified recurrent gene amplifications and a recurrent point mutation, in 16 (7.8%) and 2 (1.0%) of the urinary tract cancer patients, respectively (Fig. 1a, 1b). The c.1149G>C (*AHR*^*Q383H*^) point mutation was previously reported as an APOBEC-associated hotspot mutation in bladder cancer based on the TCGA PanCancer Atlas data ^11^. The *AHR*^*Q383H*^ point mutation was found in 11 patients out of ~11,000 patients in the TCGA PanCancer Atlas, which are mainly of primary tumor origin ^19,20^. The majority (n = 8) occurred in bladder urothelial carcinomas with the other three in hepatocellular carcinoma, papillary renal cell carcinoma, and lung adenocarcinoma. Thus, like the *AHR*^*Δe8-9*^ variant, the *AHR*^*Q383H*^ mutation is highly specific for urinary tract cancer.

AHR ligand-activated transcription factor that contains a basic Helix-Loop-Helix/PER-ARNT-SIM (bHLH/PAS) motif. AHR contains two PAS domains of which the PAS-B domain contains the ligand binding domain (LBD) (Fig. 1c) ^12,21^. AHR is part of a cytoplasmic protein complex containing HSP90, p23, and XAP2 (Fig. 1d). Upon ligand binding, AHR dissociates from the complex and translocates to the nucleus where it forms a heterodimer with ARNT ^22^. The AHR/ARNT complex subsequently binds to Xenobiotic Response Elements (XREs) in the genome to activate the transcription of target genes such as the phase I and II drug metabolizing enzymes CYP1A1 and CYP1B1 ^12^. At the protein level, the *AHR*^*Δe8-9*^ deletion results in an in-frame protein coding sequence with the loss of 84 amino acids (p. 303-387), disrupting the PAS-B domain and the C-terminal part of the ligand-binding domain of the protein. The HSP90 and XAP2 protein binding domains are predicted to be affected, while the dimerization region of the protein likely remains intact (Fig. 1c).

Among the urinary tract cancer patients in the Hartwig database, *AHR* alterations are mutually exclusive with *FGFR3* alterations and *PIK3CA* mutations, although the significance is impacted by low sample numbers (*FGFR3*: Odds ratio = 0.24 p = 0.21, *PIK3CA*: Odds ratio = 0.30 p = 0.32). The tendency of mutual exclusivity with genes from the RTK-Ras-PI3K pathway strengthens the idea that *AHR* is an independent driver of urinary tract cancer, with potential convergence on the RTK-Ras-PI3K pathway. *AHR* alterations do not show depletion with genes involved in chromatin remodeling or the tumor suppressors *TP53* and *Rb1* (Fig. 1e, Supplementary table 4).

For tumors with *AHR* amplifications, *AHR* expression is increased compared to the non-amplified urinary tract cancer samples (Fig. 1f). Urinary tract cancers with *AHR*^*Δe8-9*^ showed increased expression of AHR target genes, which was not observed in the *AHR* amplified or *AHR* non-affected samples (Fig. 1g). No RNA sequencing data was available for the 2 patients with the *AHR*^*Q383H*^ mutation.

### The *hAHR*^*Q383H*^ mutation affects ligand binding affinity and specificity

The mouse ortholog of *AHR*^*Q383H*^ (Ahr^Q377^) can form hydrogen bonds with Ahr ligands in the ligand-binding domain and mutations that change this residue affect ligand binding affinities ^23–25^. To examine if ligand binding affinities are also affected for the human AHR (hAHR), we measured the transcriptional activity of hAHR^Q383H^ and hAHR^WT^ in an XRE-luciferase reporter assay. Both the hAHR^WT^ and hAHR^Q383H^ expressing cells showed strong transcriptional activity with AHR ligands TCDD and B[a]P (Fig. 2a). Stimulation with B[a]P resulted in a higher induction of luciferase transcription for the hAHR^Q383H^ mutant than for the hAHR^WT^, suggesting a difference in ligand affinity between both variants. The AHR antagonist CH-223191 reduced transcriptional activation in hAHR^WT^ cells stimulated with TCDD or B[a]P (Fig. 2a) ^26,27^. Surprisingly, incubation of hAHR^Q383H^ with CH-223191 resulted in transcriptional activation and no antagonism was observed when CH-223191 was combined with TCDD or B[a]P (Fig. 2a). To examine this altered ligand-binding specificity, we analyzed protein localization in RPE1 cells. In the absence of exogenous ligands, both hAHR^Q383H^ and hAHR^WT^ proteins are mainly localized to the cytoplasm and both proteins show nuclear translocation upon incubation with TCDD or B[a]P (Fig. 2b, Supplementary fig. 1). Incubation with the antagonist CH-223191 also resulted in nuclear translocation of hAHR^Q383H^, while the hAHR^WT^ remains localized to the cytoplasm (Fig. 2b). The difference in responses between the hAHR^Q383H^ and hAHR^WT^ to AHR agonists and antagonists demonstrates the Q383H mutant is more sensitive to AHR pathway stimulation.

**Figure 2.**
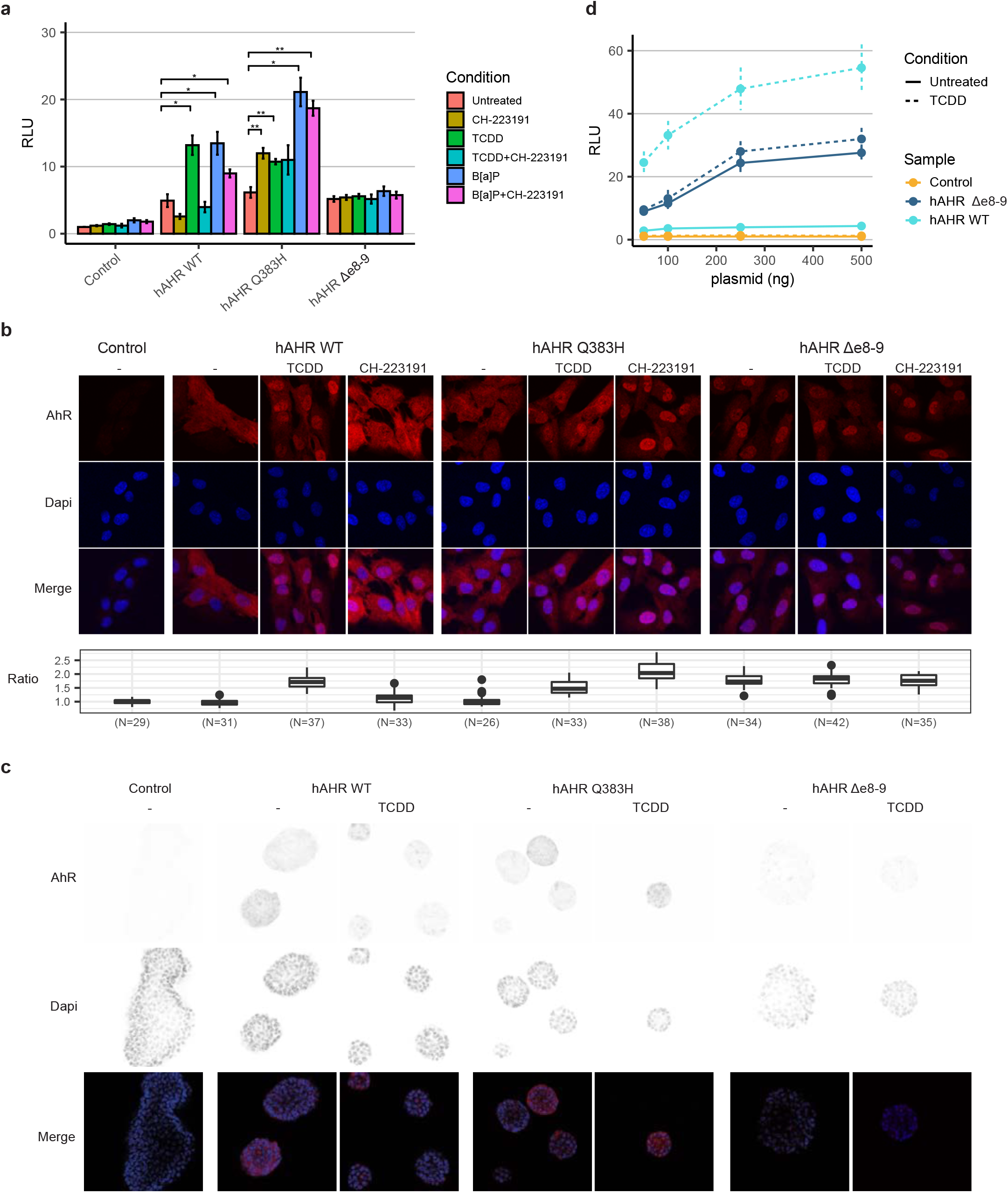
Altered protein localization and mediated transcription activation of *hAHR* mutants. a. Luciferase reporter assay of HEK293T cells transfected with empty control, *hAHR*^*WT*^, *hAHR*^*Q383H*^, or *hAHR*^*Δe8-9*^ vectors together with XRE-luciferase reporter construct. Assays were performed with different AHR ligands (TCDD and B[a]P) and AHR antagonist (CH-223191) conditions. Significance was calculated with Student’s T-test for stimulated versus untreated condition per construct. * p < 0.05, ** p < 0.01. RLU = relative luciferase unit. b. Immunofluorescence of RPE1 cells that express hAHR^WT^, hAHR^Q383H^, or hAHR^Δe8-9^ that were incubated for 4h with TCDD or CH-223191. AHR was detected with immunofluorescence (red) and DNA with Dapi (blue). The boxplots present the AHR signal ratio of nuclear to cytoplasmic signal for all individual captured cells. Untransduced cells served as negative controls (left panels). c. Immunofluorescence of mouse bladder organoids that express hAHR^WT^, hAHR^Q383H^, or hAHR^Δe8-9^ and were incubated for 24h with TCDD. AHR was detected with immunofluorescence (red) and DNA with Dapi (blue). d. Luciferase reporter assay of HEK293T cells transfected with increasing concentration of hAHR^WT^, hAHR^Δe8-9^, and empty control vectors.

### The hAHR^Δe8-9^ mutant leads to constitutive AHR pathway activation

To study the functional consequences of *AHR*^*Δe8-9*^, we created transgenic *hAHR*^*Δe8-9*^ RPE1 cells and mouse bladder organoids. In contrast with *hAHR*^*WT*^, the *hAHR*^*Δe8-9*^ protein is localized in the nucleus regardless of the presence or absence of a ligand (Fig. 2b). Expression of the *hAHR* mutants in mouse bladder organoids confirmed constitutive nuclear localization of the hAHR^Δe8-9^ mutant (Fig. 2c). In XRE-luciferase reporter assays, the degree of transcriptional activation by hAHR^Δe8-9^ was similar for all the conditions, irrespective of the presence of AHR ligands. Moreover, the AHR antagonist CH-223191 did not affect the transcriptional activation activity of hAHR^Δe8-9^ (Fig. 2a). Increased concentrations of *hAHR*^*WT*^ and *hAHR*^*Δe8-9*^ constructs in the transfections resulting in increased expression levels and revealed that hAHR^Δe8-9^ induces higher transcriptional activity levels than hAHR^WT^ in untreated conditions (Fig. 2d).

To further characterize the consequences of the constitutive nuclear localization of hAHR^Δe8-9^ we performed bulk RNA-sequencing on mouse bladder organoids expressing *hAHR*^*Δe8-9*^ or *hAHR*^*WT*^ in the presence or absence of TCDD. Untreated *hAHR*^*Δe8-9*^ organoids differentially expressed several genes compared to the *hAHR*^*WT*^ organoids, including the AHR target genes *Cyp1a1*, *Cyp1b1*, *Ahrr*, *Gsta1*, and *Tiparp* demonstrating constitutive activation of the AHR pathway (Fig. 3a, b). The expression pattern of up and down regulated genes is similar for *hAHR*^*Δe8-9*^ in untreated and TCDD treated conditions, confirming that loss of exons 8 and 9 leads to ligand-independent activation of the AHR pathway, which is in line with the upregulation of these genes in *AHR*^*Δe8-9*^ positive urinary tract cancers (Fig. 1g). Thus, *AHR*^*Δe8-9*^ leads to constitutive activation of the AHR pathway.

**Figure 3:**
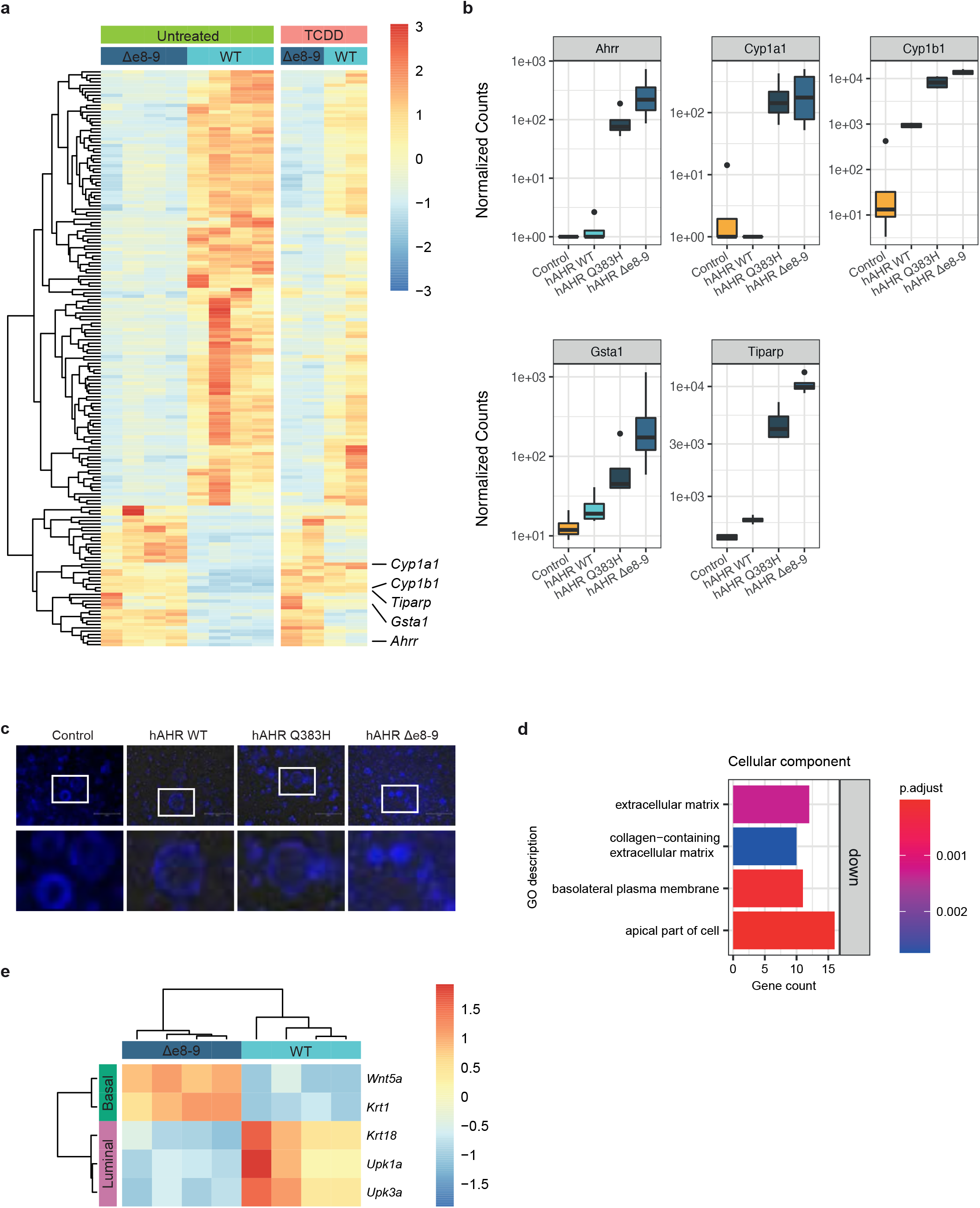
Transcriptome analysis of constitutively activated *hAHR*^*Δe8-9*^ mutant. a. Rlog values of the 172 most differentially expressed genes between the *hAHR*^*WT*^ and *hAHR*^*Δe8-9*^ expressing mouse bladder organoids in untreated and TCDD treated conditions. b. Boxplot of normalized counts of RNA reads of AHR target genes for the untreated mouse bladder organoids expressing *hAHR* mutants (n=4). c. Images of Dapi stained mouse bladder organoids expressing *hAHR* mutants. The lower panel shows the magnification of the boxed area. Scale bar 250 μm. d. GO enrichment analysis of *hAHR*^*Δe8-9*^ mutant for the cellular component domain. e. Rlog values of basal and luminal cell layer makers for *hAHR*^*Δe8-9*^ and *hAHR*^*WT*^ expressing mouse bladder organoids.

The *hAHR*^*WT*^ organoids treated with TCDD also show upregulation of the canonical AHR target genes, but do not show the same downregulated genes (Fig. 3a). This observation indicates a different effect on the transcriptome between the constitutively active hAHR^Δe8-9^ and the 24h TCDD stimulated hAHR^WT^ condition. Moreover, overexpression of *hAHR*^*WT*^ and *hAHR*^*Q383H*^ without the addition of exogenous ligands already results in modest pathway activation when compared with control organoids that are not transgenic for *hAHR* (Fig. 3b). These observations are in line with the higher induction of luciferase transcription in the untreated condition for the *hAHR*^*WT*^ and *hAHR*^*Q383H*^ constructs compared to the control in the luciferase assay (Fig. 2a).

### The *hAHR*^*Δe8-9*^ mutant induces cellular transformation

The bladder is a stratified epithelium, with stem cells that reside in the basal cell layer that support organ regeneration and renewal. Upon differentiation, the stem cells give rise to intermediate cells and luminal umbrella cells ^28^. Mouse bladder organoids enriched for undifferentiated cells have a reduced diameter of the lumen and increased thickness of the epithelial layer when compared to organoids composed of differentiated cells ^29^. We observed a mixture of cystic and compact organoids in the mouse bladder organoids expressing *hAHR*^*WT*^, indicating a heterogeneous population of differentiated and undifferentiated cells. No cystic organoids were observed in the organoids expressing *hAHR*^*Δe8-9*^ indicating a more basal stem-cell like phenotype (Fig. 3c).

We performed GO enrichment analysis on all differentially expressed genes in RNA-sequencing data of the mouse organoids to understand which processes are affected by *hAHR*^*Δe8-9*^. Most outstanding is the downregulation of genes related to the extracellular matrix and cell periphery organization (Fig. 3d, Supplementary table 5). Moreover, *hAHR*^*Δe8-9*^ expressing mouse organoids show downregulation of the differentiation markers *Upk3a*, *Upk1a*, and *Krt1B* and upregulation of the stem cell markers *Wnt5a*, and *Krt1* when compared to *hAHR*^*WT*^ organoids (Fig. 3e) ^29–31^. Together, these observations indicate that *hAHR*^*Δe8-9*^ promotes a basal stem-cell like phenotype in bladder cells.

Because RNA-seq was performed on bulk cultures, the transcriptional changes induced by *hAHR*^*Δe8-9*^ may reflect a shift in the composition of the cell types towards a more homogeneous population, or the transformation of cells towards a novel phenotype. To discriminate between these scenarios, we performed scRNA-seq on the mouse bladder organoid lines. Dimensional reduction and unsupervised clustering revealed the presence of 7 clusters (Fig. 4a). Based on the genes enriched in the different clusters, cluster 0 represents a basal phenotype (characterized by *Krt14*, *Trp63*, *Bcam* and *Agrn*), cluster 1 represents an intermediate/luminal phenotype (characterized by *Krt19*, *Krt1B*, *Upk1b*, *Cldn4*, *Cldnl, Ceacam1* and *Alcam*), and clusters 3 and 5 represent cells that are in the S-phase and M-phase of the cell cycle, respectively (Fig. 4a, 4b, Supplementary table 6). Together, these clusters represent a classical stem cell system where stem cells divide to give rise to new stem cells or to cells that differentiate. The vast majority of all cells of the control lines fall into these clusters and in cluster 4, which we were not able to link to a particular cellular phenotype. Strikingly, two clusters (clusters 2 and 6) were almost exclusively occupied by mouse bladder cells expressing *hAHR*^*Δe8-9*^ (Fig. 4a, 4c). In addition to the canonical AHR target genes such as *Cyp1b1* and *Tiparp*, cluster 2 expresses basal stem cell markers such as *Krt1l*, *Wnt5a*, *Itga6*, *Wnt4* ^29^. In addition cluster 2 is characterized by genes that are associated with urothelial cancer, such as: *Htra1* ^32^, *Cyb5r1* ^33^, *Steap1* ^33,34^, *Ptgs2* ^35,36^, and *Trib3* ^37^ (Fig. 4b). Cells in cluster 6 show upregulation of *Dsp*, *Pkp1*, *Ppl*, *Jup* and *Krt13*, which are involved in desmosome and intermediate filament cytoskeleton organization. This cluster has some overlap with markers expressed in the luminal-intermediate cluster (cluster 1) of the control cells, but with less apparent expression of the umbrella cell markers. This indicates that the cells in cluster 6 represent an intermediate cell type that however fails to differentiate towards luminal umbrella cells. Thus, the constitutive activation of the AHR pathway by *hAHR*^*Δe8-9*^ leads to a transformation of cells towards a less differentiated phenotype and the activation of genes linked to urothelial cancer.

**Figure 4:**
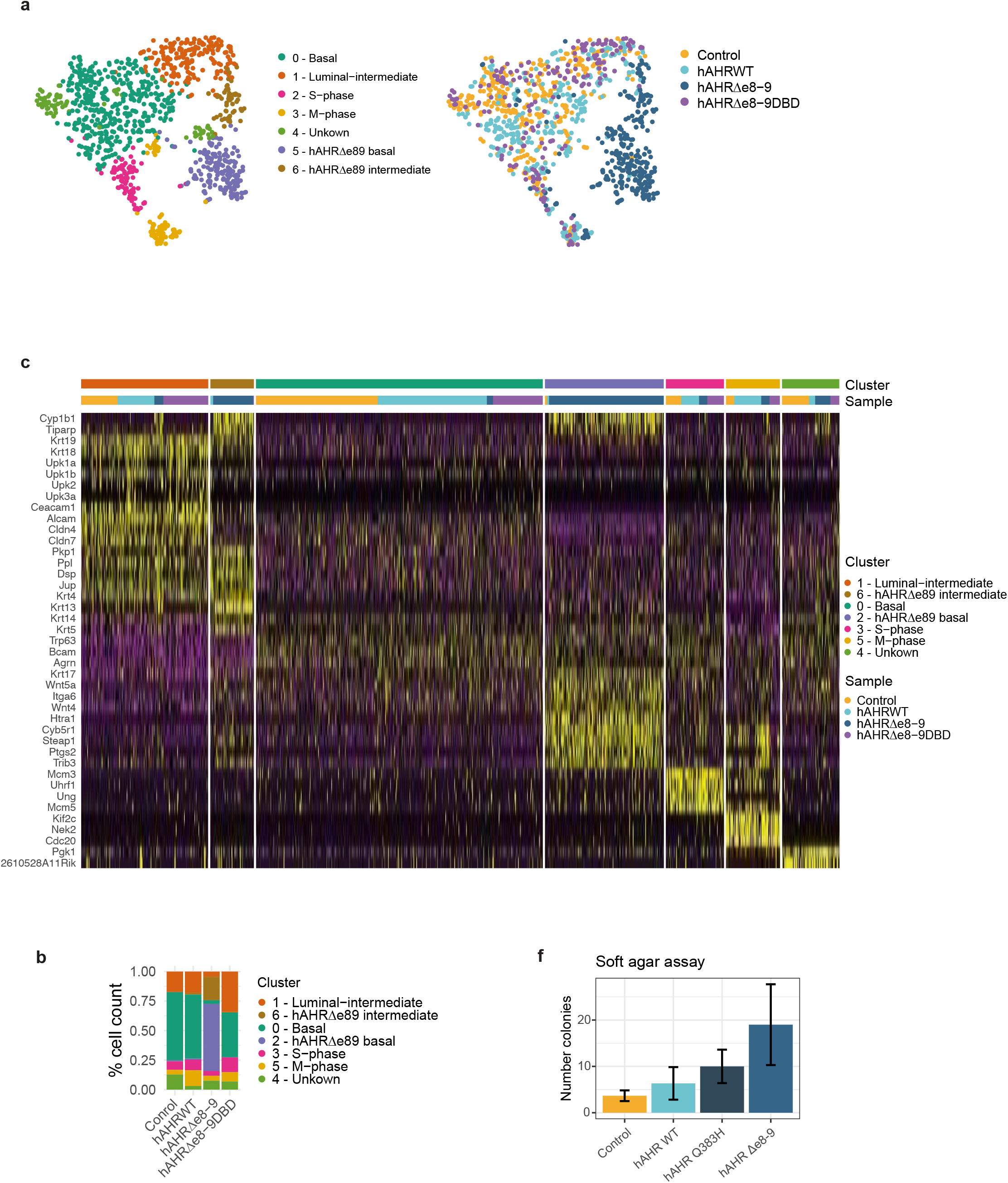
Transformed phenotype of hAHR^Δe8-9^ mutant. a. tSNE plot of single cell transcriptomics depicting clusters of the transgenic mouse bladder organoids cells. Plot is coloured for detected clusters and for the different transgenic mouse bladder organoid lines. b. Heatmap depicting expression of selected markers, which are identified by expression analysis, representative for the different identified clusters. c. Contribution of percentage of cells to the different clusters for each transgenic mouse bladder organoid line. d. Soft agar assay of mouse bladder organoids expressing hAHR mutants. Results are presented as the mean ± SD of triplicate samples.

Cells of a DNA binding deficient variant of the oncogenic *hAHR*^*Δe8-9*^ (*hAHR*^*Δe8-9DBD*^) clustered among cells of the other control lines and apart from *hAHR*^*Δe8-9*^ cells (Fig. 4a). This shows that the cellular transformation of the *hAHR*^*Δe8-9*^ organoids depends on the transcriptional activation caused by AHR and is not the result of potential transcription-independent effects caused by the deletion. To functionally explore if the transformed phenotype is accompanied by oncogenic properties, a soft agar growth assay was performed. A higher number of colonies were counted for *hAHR*^*Δe8-9*^ organoids compared to *hAHR*^*WT*^ organoids or control organoids demonstrating that *hAHR*^*Δe8-9*^ confers anchorage-independent growth to bladder cells (Fig. 4d).

Together, these observations demonstrate that transcriptional changes driven by the constitutively active *hAHR*^*Δe8-9*^ lead to a transformation of cells towards a cancerous phenotype that fails to differentiate and is able to grow independent of anchorage to the extracellular matrix.

## Discussion

In this study, we show high prevalence of *AHR* alterations in urinary tract cancers and provide functional validations to support that these aberrations are oncogenic and drive urinary tract cancer. These findings are in line with mouse studies that have shown that *AHR* overexpression or pathway activation can lead to malignant transformation of epithelial cells ^38–40^. A variety of molecular signaling pathways have been previously linked to AHR-mediated tumorigenesis whether or not in a cancer type-specific background and/or driven by AHR agonists ^13,41^. This includes the biotransformation of hydrocarbons by CYP enzymes to mutagenic intermediates which can induce DNA adducts, cross-talk of AHR with other signaling pathways, and interaction of AHR with other binding partners than ARNT to promote transcription of non-canonical genes ^42–47^. However, the underlying molecular signaling pathways driving the tumorigenesis of urinary tract cancers in patients with activating mutations in *AHR* remains unclear. The tendency of mutual exclusivity between *AHR*, *FGFR3*, and *PIK3CA* alterations may indicate convergence on the same pathway, although a fully independent parallel oncogenic pathway in bladder cancer driven by *AHR* cannot be excluded. The *hAHR*^*Q383H*^ and *hAHR*^*Δe8-9*^ variants are almost uniquely detected in urothelial cancers, which suggests this tissue is particularly sensitive to deregulated *AHR* signaling. However, our results, nor information in literature, provides clues why AHR activation is so specific for bladder cancer, so this will require further investigations.

We demonstrate that the *hAHR*^*Q383H*^ mutation leads to increased sensitivity which may also lead to overactivity of the AHR pathway in the urinary tract, thereby driving tumorigenesis. This likely also holds true for *AHR* amplifications since overexpression of *hAHR*^*WT*^ in different models shows a modest increase of AHR pathway activation compared to the controls. It is not known if the ligands that induce overactive AHR signaling in the tumors with *AHR* amplified and *AHR*^*Q383H*^ backgrounds are of environmental origin (like components in tobacco smoke) or have an endogenous source (like metabolites) ^48^. Surprisingly, we observed *hAHR*^*Q383H*^ activation upon treatment with the AHR antagonist CH-223191. This contrasts with a recent study that demonstrated reduced viability upon treatment with CH-223191 of the bladder cancer cell line KMBC2 harboring the *AHR*^*Q383H*^ mutation ^11^. A possible explanation for the apparent discrepancy in results may lie in the different types of experiments that were conducted. The reduced viability of the KMBC2 cell line after CH-223191 incubation may also be independent of alterations in AHR pathway activity.

Patients that harbor *AHR* activating mutations could potentially benefit from AHR targeted therapies ^49^. Here we show that the mode of action of AHR activation differs between the different mutations, which implies that tailored therapies depending on the underlying mutational event are required. Most classic AHR antagonists function by interference with the ligand-binding domain, but this domain is not targetable for the AHR^Δe8-9^ mutant as this domain is deleted. Therefore, functional screens to identify specific AHR^Δe8-9^ targeting compounds or the identification of essential downstream activated processes could be a next step towards the identification of novel treatment strategies for selected urinary tract cancer patients in the context of precision medicine.

## Material and Methods

### Driver gene status urinary tract cohort

Variant detection and driver likelihood status are based on the Hartwig database ^14^. Driver likelihood is introduced to select for a sample specific likelihood of each gene based on the type of variant and taking mutation load per sample into account. To select for affected genes in the urinary tract cancer samples, the driver likelihood score is set to >0.8 and detected gene fusions with a high impact are included. Visualized are the genes affected in more than 12.5% of the samples (top 12 genes). Mutually exclusivity is pairwise calculated with Fisher Exact Test based on Odd Ratio cut-offs as described in Gao et al. ^19^.

### Gene constructs

A plasmid containing the *hAHR*^*WT*^ sequence was purchased from Origene (RC209832). The Q383H point mutation was introduced with site-directed mutagenesis with forward primer cattgtaactcacagaccactaacagatg and reverse primer gttagtggtctgtgagttacaatgatataatc. The *hAHR*^*WT*^, *hAHR*^*Q383H*^, and *hAHR*^*Δe8-9*^ sequences were cloned in a lentiviral plasmid (Addgene #52961) and subsequently to pcDNA3.1 vector with the In-fusion HD Eco-dry cloning (Takarabio). Primers for lentiviral plasmid are: forward N-flag caggaccggttctaggatatcgccaccatggattacaaagacgatgacgataagaacagcagcagcgccaac, forward caggaccggttctaggatatcgccaccatgaacagcagcagcgcc, reverse ttgttgcgccggatcgcaggaatccactggatgtcaaatcag, reverse C-flag ttgttgcgccggatcgcttatcgtcatcgtctttgtaatccaggaatccactggatgtcaaatcag, forward deletion tggttgtgatgccaaagatgaggaaggaacagagca and reverse deletion gttccttcctcatctttggcatcacaaccaatagg. Primers cloning of *AHR* sequences to pcDNA3.1 vector are forward taccgagctcggatcatatcgccaccatgaacag and reverse gatatctgcagaattttacaggaatccactggatgtcaaat. For the DNA binding deficient variant of *AHR*^*Δ8-9*^, mutations were introduced to substitute the amino acids Histidine 39 and Arginine 40 with Alanines ^50^.

The luciferase reporter vector pGL3-XRE-Luc2 reporter vector was constructed using an in-house pGL3-minP-Luc2 vector containing DNA binding sequence “gggaggtacttggagcggc” with primers forward-1 gcgtgctagcccgggTTGCGTGACCctcgagatctgcgat, reverse-1 atcgcagatctcgagGGTCACGCAAcccgggctagcacgc, forward-2 aatcgataaggatccTTGCGTGACCgtcgaccgatgccct, reverse-2 agggcatcggtcgacGGTCACGCAAggatccttatcgatt.

### Cell culture

All cells were cultured in a humidified atmosphere at 37°C under 5%CO_2_ and 20%O_2_. Lentivirus particles containing *hAHR* constructs were produced by transient calcium phosphate transfection of HEK293T. Virus particles were concentrated with Lenti-X Concentrator (Takarabio) according to the manufacturer’s directions. Mouse bladder organoids were a kind gift from drs Hans Clevers and Jasper Mullenders and were cultured as previously reported^51^. In short, mouse bladder organoids were plated in droplets of ice-cold Matrigel (Corning). After the matrigel had solidified, a complete mouse bladder medium was added consisting of Advanced DMEM/F-12 supplemented with HEPES, Glutamax, 1% penicillin, 1% streptomycin, Nicotinamide (10 mM), N-Acetylcysteine (1.25 mM), FGF10 (100 ng/mL of Peprotech 100-26), FGF7 (25 ng/mL of Peprotech 100-19), A83-01 (500 nM), B27 (ThermoFisher 17504001) and primocin (Invivogen). Organoids were split weekly either through mechanical shearing with a pulled Pasteur’s pipet or by dissociation using TrypLE. To increase survival the medium was supplemented with ROCK inhibitor (Y-27632, 10 μM) after splitting.

RPE1 and HEK293T cells were cultured in DMEM, 10% fetal bovine serum, 1% penicillin, and 1% streptomycin. After transduction, RPE1 cells and the mouse bladder organoids were placed on 10-15 μg/ml and 1 μg/ml puromycin selection, respectively. Cells were incubated with 10 nM TCDD (LGC), 10 μM B[a]P (sigma), or 10 μM CH-223191 (Sanbio) or vehicle alone (DMSO) for 24h unless otherwise stated.

### Immunofluorescence

RPE1 cells expressing *hAHR* mutants were plated on coverslips and incubated with TCDD, B[a]P, CH-223191 or untreated (DMSO) for 4 or 24h. Cells were fixed in 4% formaldehyde, permeabilized with 0.3% Triton X-100 in PBS, and blocked with 4% BSA in PBS. AHR was stained with Anti-AHR antibody (Abcam ab190797) in blocking buffer. After incubation with the primary antibody, cells were washed followed by incubation with secondary antibodies conjugated with Alexa fluor dyes. Subsequently, the cells were washed, counterstained with DAPI, and mounted on microscope slides. Images were taken with predefined settings on Zeiss LSM510 microscope and analyzed with Fuij/imageJ. Ratios of AHR signal intensities were calculated by determining average grey value intensities for each nucleus, based on dapi staining, and for each corresponding cytoplasm, as 1μM band around each nucleus.

The first steps in the immunofluorescence procedure for the mouse bladder organoids are different from the procedure described previously. First, dispase was added to a final concentration of 1mg/ml to a warm medium and incubated for 30 min at 37°C to dissolve the Matrigel. Organoids were harvested by gentle pipetting, collected in a tube, and washed with cold Advanced DMEM/F12. The organoids were centrifuged, washed in PBS followed by another centrifugation step. The resulting organoid pellet was resuspended in 100μl PBS and transferred to Lab-Tek Chamber slides. The organoids were fixed by the addition of 4% formaldehyde followed by 1h incubation at RT. The organoids attached to the bottom of the chamber slide were washed twice with PBS. Subsequently, the organoids were incubated in blocking buffer and the immunofluorescence procedure was followed as described for the RPE1 cells.

### Luciferase reporter gene assay

HEK293T cells were cultured in 24-wells plates and transiently transfected with pcDNA3.1 empty vector or pcDNA3.1 vector containing the expression cassette of *hAHR*^*WT*^, *hAHR*^*Q383H*^, or *hAHR*^*Δe8-9*^ (10 ng), together with the pGL3-XRE-Luc2 reporter vector (1000 ng) and pRL-CMV Renilla vector (2 ng). 24h after transfection, cells were treated with ligand TCDD (50 nM) or B[a]P and/or antagonist CH223191 for approximately 24h. Luciferase activity was measured with the Dual-Luciferase reporter assay system and normalized for Renilla activity (Promega, Madison, WI). The results are averages of at least three independent experiments assayed in duplicate ± SEM. Results are presented as change relative to the empty vector in the untreated (DMSO) condition.

### RNA-sequencing

The RNA from the transgenic mouse bladder organoids was isolated with Trizol (ThermoFisher) according to the manufacturer’s instructions. RNA-seq libraries were prepared using TruSeq Stranded Total RNA Library Prep Kit (Illumina) according to the manufacturer’s protocol. RNA-seq libraries were pooled and sequenced on a NextSeq2000 (Illumina) in 1×50bp single end mode. RNA sequencing reads were aligned against mouse reference genome GRCm38 using STAR and the number of reads mapping to genes was counted using featureCounts all by using a custom in-house pipeline (https://github.com/UMCUGenetics/RNASeq-NF). The Bioconductor package DESeq2 was used to normalize raw read counts and to perform differential gene expression analysis with apeglm shrinkage ^52,53^. The analyses were performed with significant (P.adjust<0.05) and differentially expressed (Log2FoldChange>2.5) genes with exception of luminal and basal marker analyses where smaller differences were included (Log2FoldChange>1.5). The Bioconductor package clusterProfiler and Revigo were used for GO enrichment analysis^54,55^.

### scRNA-seq

The transgenic mouse bladder organoids were cultured in 2D and dissociated to single cells using a 10 min incubation with TryLE. Cells were sorted into 384-well capture plates and the scRNA library preparation and sequencing were performed according to the SORT-seq protocol by Single Cell Discoveries B.V. ^56^. For all single cells, reads were aligned to the mouse reference genome GRCm38 and Sort-seq read counts were filtered to exclude reads with identical library-, cell-and molecules ^56^. With the Seurat R package, low quality cells were removed by a cut-off of 10000 transcripts per cell and the data was normalized and scaled ^57^. The top 2000 most variable genes in the dataset were identified and used for principal component analysis to determine dimensionality and clustering of the dataset. Cluster gene markers were detected using a Wilcoxon rank sum test between each cluster and the rest of the cells in the dataset with a bonferroni correction for multiple testing.

### Soft agar assay

3% agarose (REF11388991001) was dissolved and autoclaved in 100 ml EBSS. One volume of melted 3% agarose was mixed with four volumes of Advanced DMEM/F12 to obtain a 0.6% solution. This mixture was added to 6-well plates (1.5 ml/well) in which the gels were allowed to solidify. Subsequently, Tryple was used to prepare single-cell suspensions of the mouse bladder organoids. The cells were counted and for each condition, 2.5 ml of cell suspension was prepared at a concentration of 1×10^e4^ cells/ml in complete medium. The cell suspension was mixed 1:1 with a warm 0.6% agarose solution to get 0.5 ×10^e4^ cells/ml in 0.3% agarose solution. Per well 1.5 ml agarose/cell mixture was plated. The next day 300 μl of the medium was added and the cells were refed every 2-4 days. After 3 weeks the cells were stained with nitroblue tetrazolium chloride solution and pictures were made. Colonies were counted using ImageJ.

## Acknowledgments

We are grateful to Jasper Mullenders and Hans Clevers for providing the mouse bladder organoids. We thank Livio Kleij for his support with microscopy, Single Cell Discoveries for support with single cell RNA sequencing, the Utrecht Sequencing Facility for providing sequencing service, and the UMC Utrecht Bioinformatics Expertise Core for data analysis and data handling. The Utrecht Sequencing Facility is subsidized by the University Medical Center Utrecht, Hubrecht Institute, Utrecht University, and The Netherlands X-omics Initiative (NWO project 184.034.019). The UMC Utrecht Bioinformatics Expertise Core is subsidized by the University Medical Center Utrecht, Center for Molecular Medicine. This publication and the underlying study have been made possible partly based on the data that Hartwig Medical Foundation and the Center of Personalised Cancer Treatment (CPCT) have made available to the study.

## Data availability

The bulk RNA-sequencing and the scRNA-seq data of the mouse bladder organoids have been deposited in ENA with the accession code PRJEB49233.

## Author contributions

JMV, E Kuijk, NB, DW and EC designed and performed experiments including data analysis and interpretation. AB, and E Kalkhoven designed and performed the luciferase assay experiments. CS and PP conducted Hartwig database analysis. BMF provided Weill Cornell Medicine data analysis. PP and EC conceived the original idea. EC and E Kuijk supervised the project. JMV, E Kuijk, and EC wrote the manuscript with input from all authors.

## Supplementary information

Supplementary table 1: Overview urinary tract cancer sample data from Hartwig database

Supplementary table 2: Overview AHR^Δe8-9^ breakpoint information

Supplementary table 3: AHR RNA-sequencing splice junction detection in urinary tract cancer patients from Weill Cornell Medicine

Supplementary table 4: Driver gene status in urinary tract cancer samples based on Hartwig database.

Supplementary table 5: GO enrichment analysis details of mouse bladder organoids expressing hAHR^Δe8-9^

Supplementary table 6: Gene markers per cluster in scRNA-seq data of the mouse bladder organoids

Supplementary figure 1: AHR protein localization after incubation with B[a]P or CH-223191. Immunofluorescence of RPE1 cells that express hAHR^WT^, hAHR^Q383H^, or hAHR^Δe8-9^ that were incubated for 4h with B[a]P or with AHR antagonist CH-223191. AHR was detected with immunofluorescence (red) and DNA with Dapi (blue). The boxplots present the AHR signal ratio of nuclear to cytoplasmic signal for all individual captured cells. Untransduced cells served as negative controls (left panels).

