## Supplementary figures and images for "Recurrent exon-deleting activating mutations in *AHR* act as drivers of urinary tract cancer"

### Supplementary Figure 1

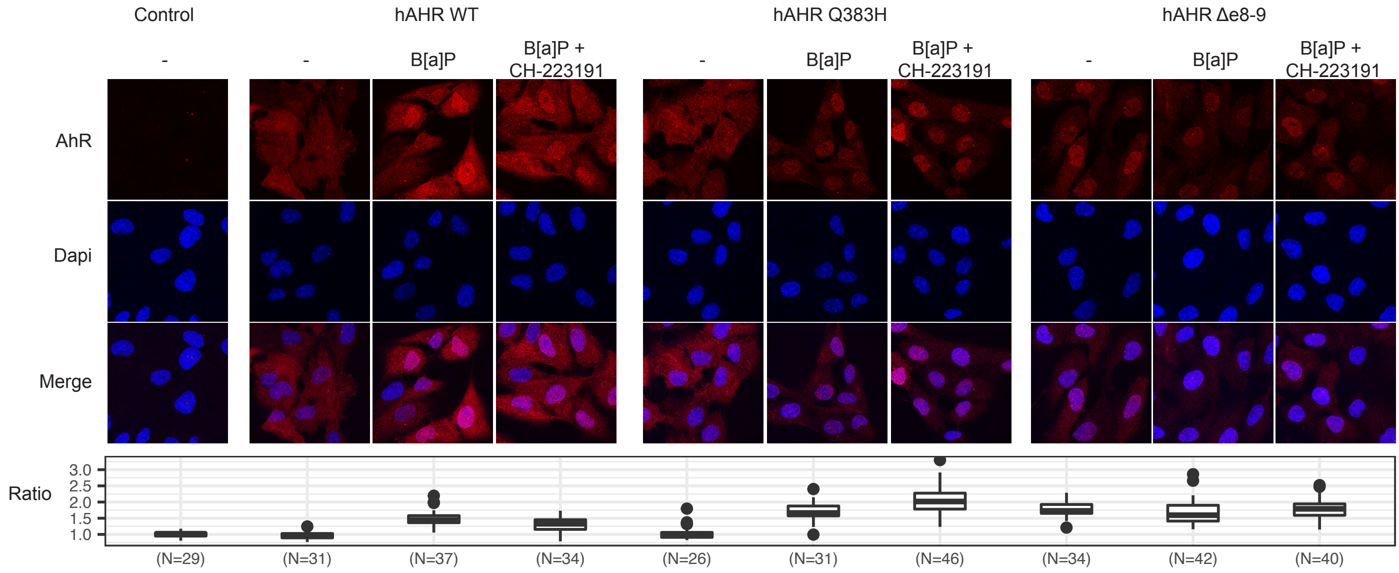
